# Efficient generation of CAR NK cells from human umbilical cord blood CD34^+^ stem and progenitors for democratizing affordable immunotherapy

**DOI:** 10.1101/2024.07.30.605741

**Authors:** Jianhuan Li, Yao Wang, Xiujuan Zheng, Yunqing Lin, Qitong Weng, Xiaofei Liu, Yang Geng, Hongling Wu, Lijuan Liu, Huan Peng, Bingyan Wu, Dehao Huang, Chengxiang Xia, Tongjie Wang, Mengyun Zhang, Xin Du, Hui Zeng, Fang Dong, Yingchi Zhang, Xiaofan Zhu, Fangxiao Hu, Jinyong Wang

## Abstract

Chimeric antigen receptor (CAR) natural killer cells (CAR NK) cells, leveraging safety and not requiring HLA match in adoptive infusion, have emerged as promising alternative cells to CAR-T cells for immunotherapies. High and multiple doses of CAR NK cell infusions are essential to maintain therapeutic efficacy in clinical trials. This requires efficient methods for generating large-scale CAR NK cells and significantly reducing CAR engineering costs. In this study, we develop a three-step strategy to generate highly high yields of induced NK (iNK) and CAR iNK cells from human umbilical cord blood CD34^+^ hematopoietic stem and progenitor cells (CD34^+^ HSPCs). Starting from a single umbilical cord blood CD34^+^ HSPC, our reliable method efficiently produces 14-83 million mature iNK cells or 7-32 million CAR iNK cells with high expression levels of CD16 and zero T cell contaminations. Introducing CAR expression elements at the HSPC level reduces the quantities of CAR pseudoviruses to 1 / 140.000 - 1 / 600,000 compared to engineering CARs in mature NK cells. The iNK and CAR iNK cells, including fresh cells and thawed cells from cryopreserved conditions, demonstrate remarkable tumoricidal activities against various human cancer cells and significantly prolong the survival of human tumor-bearing animals. The high yields of CAR NK cells and negligible costs of CAR engineering of our method support the broad applications of CAR NK cells for treating cancer patients.

## Introduction

Cellular immunotherapy, particularly chimeric antigen receptor (CAR) T cell therapy, has revolutionized the treatment of B cell malignancies. Despite the remarkable efficacy of CAR-T cells, their high costs, severe toxicities, and limited cell sources have spurred interest in developing NK cells as an alternative to universal cellular immunotherapy^1–3^. NK cells have the inherent ability to directly kill neoplastic, virally infected, and certain stressed cells due to abnormal expression of NK cell ligands such as MHC class I and class I-like molecules^4^. NK cells possess unique advantages of low side effects, such as GvHD and CRS observed in CAR-T cell therapy. CAR NK cell therapy is promising in achieving dual targeting and universal nonspecific killing effects in the treatment of tumors with low health risks^5^.

Human NK cells have a physiological turnover time of about two weeks in blood circulation^6^. The persistence of allogeneic NK cells in patients ranges from a few days to a few weeks, with an average of 7 days in most clinical trials^7^. New methods are needed to generate abundant CAR NK cells and reduce CAR engineering costs, considering the high dose (107 / kg) and multiple dose requirement for CAR NK cell therapy in clinical trials to enhance efficacies^8, 9^.

CD34^+^ HSPCs can differentiate into multilineage blood and immune cells, including NK cells. A cytokine-based culture system has produced mean output yields of 1,879-4,450 NK cells from a single HSPC. However, at the end of the culture, HSPC-NK cells express deficient levels of CD16 protein (3.0% ± 2.4)^10^, indicating low ADCC activity in the engagement of adaptive immune cells^11^. Direct transplantation of engineered CAR HSPCs results in the generation of CAR NK cells *in vivo*, with derivations of other lineage CAR cells, including CAR T, CAR Mye, and CAR B cells^12^, which brings unknown health risks.

CD34+ HSPC culture and expansion techniques have advanced over the past decades^13, 14^. Artificial organoid aggregates combining seed cells with feeder cells significantly increase the generated efficiencies of induced T cells and NK cells from stem and progenitor cells^15, 16^ Therefore, integrating HSPC expansion and organoid induction can further promote the efficiency of NK cell generation from CD34^+^ HSPCs.

This study introduces a comprehensive method to potentially derive trillions of iNK and CAR iNK cells from a single umbilical cord blood unit of CD34^+^ HSPCs. This method includes three steps: CD34^+^ HSPC expansion (Day 0-Day 14), NK lineage through organoid aggregates (Day 14-Day 28), and maturation of NK cells and large-scale proliferation in cell culture bags (Day 28-Day 49). Interestingly, a single CD34^+^ cell ultimately produced 1.4 × 10^7^ ± 0.1 × 10^7^ iNK cells and 7.6 × 10^6^ ± 1.2 × 10^6^ CD19-CAR iNK cells on day 42, and much higher yields of iNK cells (8.3 × 10^7^ ± 0.7 × 10^7^) and CD19-CAR iNK cells (3.2 × 10^7^ ± 0.2 × 10^7^) on day 49 using our technique, which significantly increased iNK cell generation efficiency to 6,918-20,224 times over the previous method. In calculation, a single umbilical cord blood unit of CD34^+^ HSPCs has the potential to produce trillions of CD19-CAR iNK cells using our technique. Interestingly, our method sharply reduces the amounts of CAR pseudoviruses to 1/140,000-1.600,000 compared to the conventional approaches of engineering CARs into mature NK cells. NK cells and CD19-CAR iNK cells, including thawed cells from cryopreserved conditions, show ideal immune activities and solid tumor eradication abilities for six months. Our method strongly supports the translational prospects for democratizing affordable CAR iNK cell therapy for the general treatment of patients.

## Results

### Efficient generation of iNK cells and CD19-CAR iNK cells from CD34^+^ HSPCs

We developed a three-step culture system to generate trillions of human iNK and CAR iNK cells from a single umbilical cord blood unit of CD34^+^ HSPCs. This system integrates strategies of HSPC expansion (Step I), organoid induction (Step II), and NK cell maturation and proliferation (Step III) (Fig. 1a). To generate iNK CAR cells with meager cost of CAR engineering, we introduced CAR expression elements in the primary CD34^+^ HSPC stage before any expansion, rather than in the mature NK stage. CD34 + HSPCs were initially stimulated for 48 hours and then transduced with CD19-CAR pseudoviruses at multiplicity of infection (MOI) 10 through spin infection. Twelve hours after infection, the transduced CD34^+^ HSPCs were immediately used for expansion culture. To enormously expand CD34^+^ HSPC and CD19-CAR CD34^+^ HSPC (CD19-CAR HSPC), we established an expansion system using AFT024 cells as feeders and HSC culture formula^17–19^. In detail, we seeded AFT024 cells (1 × 10^5^ cells/well, 20 wells) in a 24-well plate, followed by irradiation (20 Gy) to arrest feeder growth. CD34^+^ HSPCs or CD19-CAR HSPCs (5 × 10^4^ cells/well, 20 wells) were seeded in irradiated AFT024 cells for the first round of 7-day expansion in the presence of HSC culture medium^14, 20^. After the first round of expansion, all cells were collected and seeded equally into six flasks (1 × 10^7^ cells/flask) containing irradiated AFT024 cells for the second round of 7-day expansion in the presence of HSC culture medium. To efficiently drive differentiation of the NK lineage from CD34^+^ HSPC, we used the organoid aggregate induction method we established previously^16^. We combined 5 × 10^5^ expanded CD34^+^ HSPCs or CD19-CAR HSPCs with 2.5 × 10^7^ OP9 cells to form 50 organoids (1 × 10^4^ expanded CD34^+^ HSPCs or CD19-CAR HSPCs and 5 × 10^5^ OP9 cells per organoid aggregate). These organoids were seeded in transwells followed by a 14-day induction of the NK lineage. On day 28, all organoids on the membrane were digested into single cells, which were immediately transferred to a 1 L cell culture bag (50 organoid-derived single CD45^+^ cells derived from organoids per bag, approximately 1 × 10^8^ cells/bag) for a 7-day culture to achieve maturation and proliferation of iNK cells. On day 35, all cells in the cell culture bag were harvested and distributed into multiple cell culture bags (1 × 10^8^ cells/bag) for another seven days of proliferation of iNK cells or CAR iNK cells.

**Fig.1.**
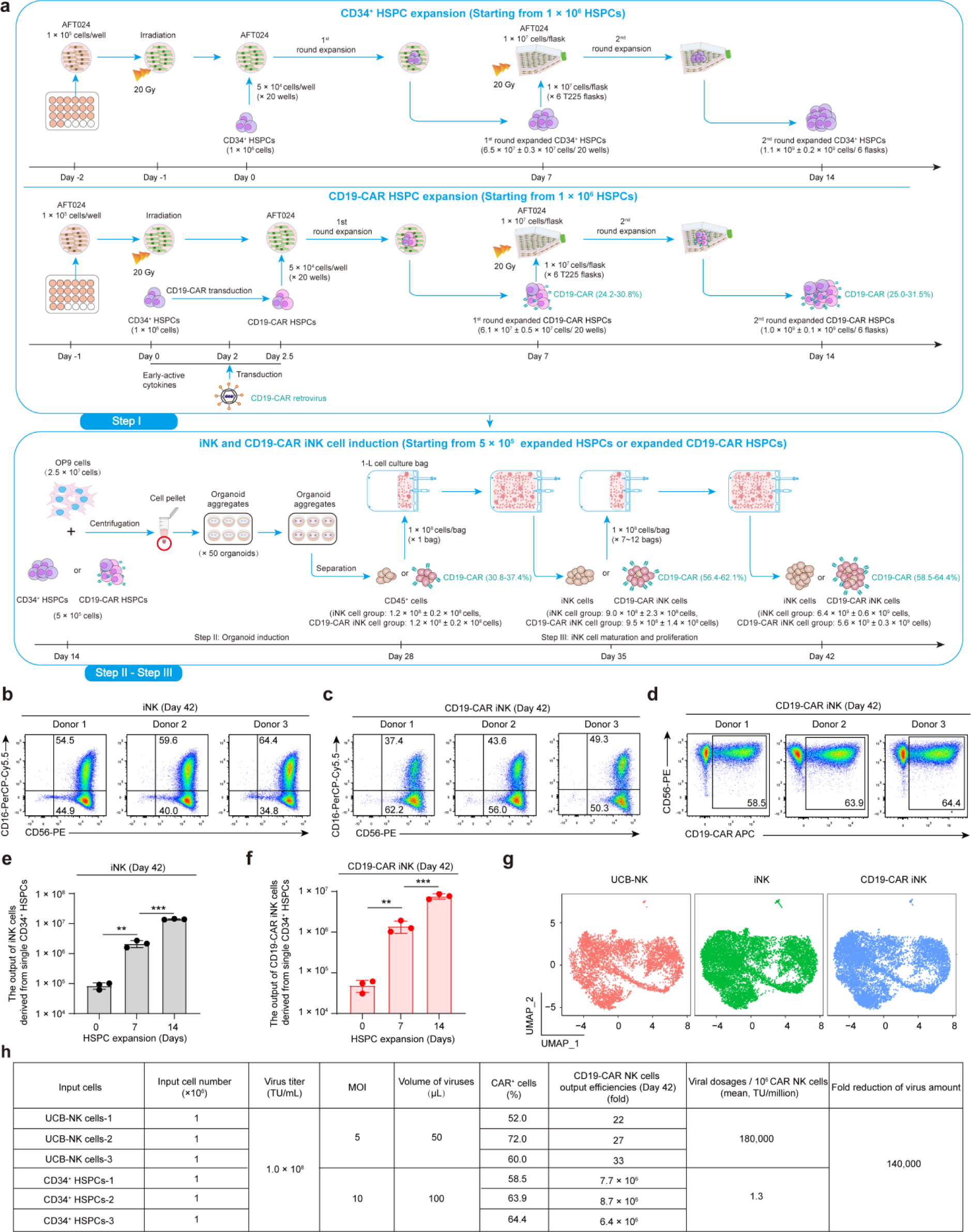
A three-step strategy for generating iNK cells and CD19-CAR iNK cells from CD34^+^ HSPCs. **a**, Schematic diagram showing the generation of iNK cells and CD19-CAR iNK cells from CD34^+^ HSPCs or CD19-CAR HSPCs. Step I was for the expansion of CD34^+^ HSPC or CD19-CAR CD34^+^ HSPC. AFT024 cells (1 × 10^5^ cells /well) were seeded in the 24-well plate and then irradiated (20 Gy). CD34^+^ HSPCs or CD19-CAR HSPCs (5 × 10^4^ cells/well) were seeded on the irradiated AFT024 cells and cocultured for the 1^st^ round. On day 7, CD34^+^ HSPC or CD19-CAR HSPC from the first round expansion were collected and seeded in the new irradiated AFT024 cells (1 × 10^7^ cells /flasks) for the second round expansion. Step II was for the differentiation of the NK lineage using the organoid aggregate induction method. CD34^+^ HSPCs or CD19-CAR HSPCs (1 × 10^4^ cells/organoid) collected from Step I were combined with OP9 cells (5 × 10^5^ cells/organoid) to prepare organoid aggregates, which were seeded in transwell and cultured for 14 days for induction of iNK cells or CD19-CAR iNK cells. Step III was for the maturation and proliferation of NK cells. CD45^+^ cells collected from Step II were transferred to cell culture bags and cultured for 14 days. **b-c,** flow cytometric analysis of the immune phenotypes of iNK cells (**b**) and CD19-CAR iNK cells (**c**) (CD56+ CD16^+/-^). Data were collected from three donor umbilical cord blood units. **d,** flow cytometric analysis of CD19-CAR expression (CD56^+^CD19-CAR^+^). **e-f,** Statistical analysis of the output efficiencies of iNK cells (CD45^+^CD3^-^CD56^+^CD16^+/-^) or CD19-CAR iNK cells (CD45^+^CD3^-^CD56^+^CD16^+/-^CD19-CAR^+^) derived from single CD34^+^ HSPCs or CD19-CAR HSPCs. Day 0, fresh CD34^+^ HSPCs. Day 7, 7-day expanded CD34^+^ HSPCs. Day 14, 14-day expanded CD34^+^ HSPCs. **g,** UMAP visualization of UCB-NK cells, iNK cells, and CD19-CAR iNK cells. **h,** Table showing the calculated quantities of CD19-CAR retroviruses (TU). The viral particles required to generate 1.0 × 10^6^ CD19-CAR iNK cells and CD19-CAR NK cells were compared. Data were presented as means ± SD. Independent two-tailed t test (**e** and **f**). NS, not significant, \*\**P* < 0.01, \*\*\**P* < 0.001.

With the three-step culture system in the absence of conventional NK cell expansion feeders^21^, we successfully harvested mature iNK cells and CD19-CAR iNK cells on day 42 with more than 99.0% purity (CD45^+^CD56^+^CD16^+/-^) and high expression levels of CD16 (37.4-64.4%) signaled the functional maturation of NK of possessing antibody-dependent cytotoxicity (Fig. 1b, c)^16^. The ratios of CD19-CAR positive iNK cells derived from three independent donor CD34^+^ HSPCs after CAR engineering were 58.5%, 63.9% and 64.4% respectively, with an average of 62.3% ± 3.3% (mean ± SD) (Fig. 1d). In calculation, single CD34^+^ HSPCs without expansion (Day 0) produce 8.5 × 10^4^ ± 2.1 × 10^4^ iNK cells and 4.9 × 10^4^ ± 1.6 × 10^4^ CD19-CAR iNK cells (mean ± SD, n=3). Single CD34^+^ HSPCs that undergo 7-day expansion (Day 7) produce 2.1 × 10^6^ ± 0.6 × 10^6^ iNK cells and 1.4 × 10^6^ ± 0.5 × 10^6^ CD19-CAR iNK cells (mean ± SD, n=3). Furthermore, single CD34^+^ HSPCs undergoing 14-day expansion (Day 14) produce 1.4 × 10^7^ ± 0.1 × 10^7^ iNK cells and 7.6 × 10^6^ ± 1.2 × 10^6^ CD19-CAR iNK cells (mean ± SD, n=3) (Day 7 vs. Day 0, *P* < 0.01, Day 14 vs. Day 7, *P* < 0.001) (Fig. 1e, f). A 14-day CD34^+^ cell expansion step resulted in eventual output efficiencies of iNK cells and CD19-CAR iNK cells more than 160 times compared to fresh HSPCs. Single cell RNA-seq analysis demonstrated that most iNK cells and CD19-CAR iNK cells projected well to activated NK cells from human umbilical cord blood (Fig. 1g). To prepare 1 million CAR NK cells, our method significantly reduced the quantities of CAR pseudoviruses to 1 / 140,000 compared to conventional techniques for engineering CARs into mature NK cells (Fig. 1h)^22^. Furthermore, iNK and CD19-CAR iNK cells could expand slightly in a bag-based culture system for another seven days. We harvested iNK cells (Supplementary Fig. 1a) and CD19-CAR iNK cells (Supplementary Fig. 1b) on day 49, which exhibited further increases in CD16 expression levels (iNK cell group, 67.8% ± 3.4%, CD19-CAR iNK cell group, 53.5% ± 6.2%, mean ± SD, n = 3 in each group), stable expression ratios of CD19-CAR in CD19-CAR iNK cells (62.9% ± 5.7%, mean ± SD, n = 3) (Supplementary Fig. 1c), and much higher yields of iNK cells (8.3 × 10^7^ ± 0.7 × 10^7^, mean ± SD, n = 3) (Supplementary Fig. 1d) and CD19-CAR iNK cells (3.2 × 10^7^ ± 0.2 × 10^7^, mean ± SD, n = 3) (Supplementary Fig. 1e). 21-day bag-based culture further reduced the quantity of quantities of CAR pseudovirus to 1/600,000 when compared to conventional methods of engineering CARs into mature NK cells (Supplementary Fig. 1f).

We monitored the immune phenotypes and summarized the related ratios and cell numbers throughout the process. We first analyzed the ratios of CD34^+^ cells during the expansion step. On day 7, 92.3% ± 3.6% (HSPC group, mean ± SD, n=3) cells retained CD45^+^CD34^+^ HSPC phenotypes. Even with engineered CAR expression elements, 86.0% ± 2.3% (CD19-CAR HSPC group, mean ± SD, n=3) cells still maintained the CD45^+^CD34^+^ HSPC phenotypes (Fig. 2a). In comparison, we did not observe significant differences in CD34 expression ratios between the HSPC group and the CD19-CAR HSPC group after seven days of expansion (*P* > 0.05). However, these ratios decreased to 71.9% ± 4.7% (HSPC group, mean ± SD, n=3) and 61.0% ± 2.4% (CD19-CAR HSPC group, mean ± SD, n=3) on day 14, respectively (Fig. 2b). It is noteworthy that the expression of CD19-CAR slightly decreased CD34 positive cell rates after 14 days of expansion (HSPC group vs. HSPC group CD19-CAR, *P* < 0.05) (Fig. 2b). Interestingly, 1 × 10^6^ CD34^+^ HSPC cells resulted in 6.4 × 10^7^ ± 0.3 × 10^7^ (CD34^+^ HSPC group) and 6.1 × 10^7^ ± 0.5 × 10^7^ (CD19-CAR HSPC group) CD34^+^ cells on day 7. A 14-day expansion further yielded 1.1 × 10^9^ ± 0.2 × 10^9^ (CD34^+^ HSPC group) and 1.0 × 10^9^ ± 0.1 × 10^9^ (CD19-CAR HSPC group) CD34^+^ cells, respectively (mean ± SD, n=3) (Fig. 1a and Fig. 2c).

**Fig.2.**
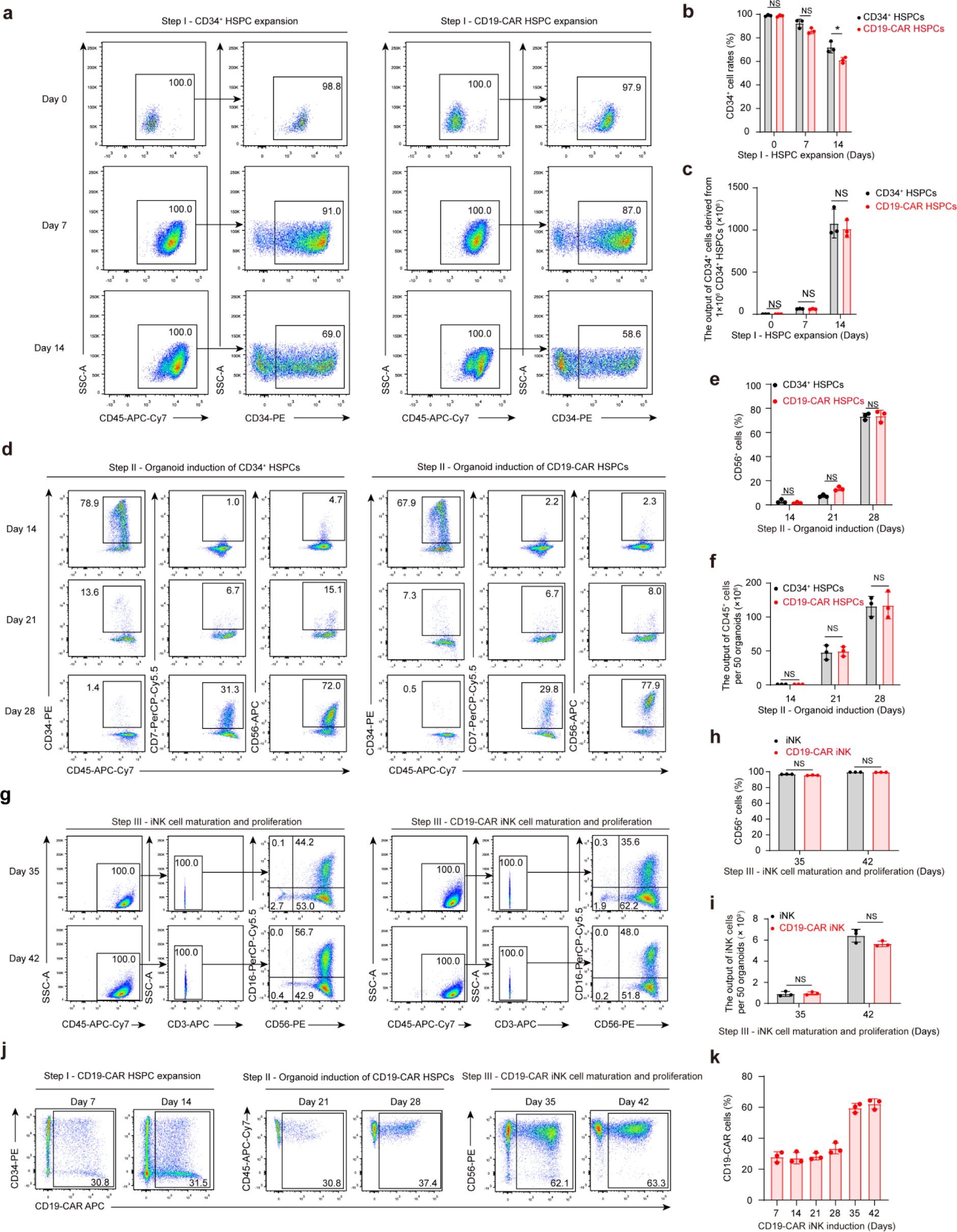
Cellular kinetics in the three-step strategy to generate iNK cells and CD19-CAR iNK cells. **a**, Representative FACS plots showing the ratios of CD34^+^ HSPCs (CD45^+^CD34^+^) during expansion (Step I) of CD34^+^ HSPCs (Left panel) and CD19-CAR HSPCs (CD45^+^CD34^+^) (Right panel) (Day 0, Day 7, and Day 14). **b,** Statistical analysis of the ratios of CD34^+^ HSPCs and CD19-CAR HSPCs (CD45^+^CD34^+^) during step I. (n = 3 in each group, n indicated three donor umbilical cord blood units). **c,** Statistical analysis of the output of CD34^+^ HSPCs and CD19-CAR HSPCs derived from 1.0 × 10^6^ CD34^+^ HSPCs in Step I. (n = 3 in each group, n indicated three donor umbilical cord blood units). **d,** Representative FACS plots showing the ratios of CD45^+^CD34^+^, CD45^+^CD7^+^, and CD45^+^CD56^+^ cells of the organoid aggregates derived from CD34^+^ HSPCs and CD19-CAR CD34^+^ HSPCs in Step II. **e,** Statistical analysis of the ratios of CD45^+^CD56^+^ cells in step II. (n = 3 in each group, n indicated three donor umbilical cord blood units). **f,** Statistic analysis of CD45^+^ cell outputs derived from 50 organoids in Step II. (n = 3 in each group, n indicated three donor umbilical cord blood units). **g,** Representative FACS plots showing the phenotypes (CD45^+^CD3^-^CD56^+^CD16^+/-^) of total cells on day 35 and day 42 during cell maturation and proliferation of iNK (Left panel) or CD19-CAR iNK (Right panel) cell maturation and proliferation (Step III). **h,** Statistical analysis of the ratios of CD45^+^CD3^-^CD56^+^cells on day 35 and day 42 in Step III. (n = 3 in each group, n indicated three donor umbilical cord blood units). **i,** Statistical analysis of the output of CD45^+^CD3^-^CD56^+^cells derived from 50 organoids on day 35 and day 42 in step III. (n = 3 in each group, n indicated three donor umbilical cord blood units). **j,** Representative FACS plots showing the expression of CD19-CAR of total cells at the indicated time points. **k,** Statistical analysis of the ratios of CD45^+^CD19-CAR^+^cells at the indicated time points. (n = 3 in each group, n indicated three donor umbilical cord blood units). Data were presented as means ± SD. Independent two-tailed t test (**b**, **c**, **e**, **f, h**, and **i**). NS, not significant, \**P* < 0.05.

To increase the efficiency of NK cell generation, we drove expanded CD34^+^ HSPC and CD19-CAR HSPC toward the NK cell lineage via an efficient organoid induction approach^16^. From day 14 to day 21, the CD34^+^ population decreased, while the proportion of cells in the NK lineage that express CD7 and CD56 gradually increased (Fig. 2d, e). On average, 50 organoids produced 1.2 × 10^8^ ± 0.2 × 10^8^ CD45^+^ cells (CD34^+^ HSPC group) and 1.2 × 10^8^ ± 0.2 × 10^8^ (CD19-CAR HSPC group) CD45^+^ cells, including precursor cells and NK cells, after 2-week organoid induction (Fig. 1a and Fig. 2f).

Subsequently, every 50 organoids as a group were digested in single cell suspensions followed by transferring them to commercial 1-L cell culture bags for 14-day NK cell maturation and proliferation^16^. After seven days of bag culture, the NK precursor cells matured into the NK cell phenotype (CD45^+^CD3^-^CD56^+^), which reached 97.0% ± 0.2% (iNK cell, mean ± SD, n=3) and 95.6% ± 0.4% (CD19-CAR iNK cell, mean ± SD, n=3) of total CD45^+^ cells. After 14-day bag culture, NK cell purity reached 99.6% ± 0.0% (iNK cell group, mean ± SD, n=3) and 99.4% ± 0.1% (CD19-CAR iNK cell group, mean ± SD, n=3). (Fig. 2g, h). On average, single cells of 50 organoids as input produced 6.4 × 10^9^ ± 0.6 × 10^9^ (iNK cell group) and 5.6 × 10^9^ ± 0.3 × 10^9^ (CD19-CAR iNK cell group) CD56^+^ iNK cells after two weeks of maturation and proliferation of NK cells (Fig. 1a and Fig. 2i). To maintain the optimal activity of NK cells, we collected iNK cells and CD19-CAR iNK cells on day 42.

Unlike the traditional NK cell expansion method using human tissue-derived NK cells as input, bringing risks of T cell contamination, our method produced over 99.0% pure NK cells with zero T cell contaminations (Fig. 2g). Notably, during the bag culture period, we observed significant increases in CD16 positive iNK cells, rising from 44.2% to 56.7% (iNK cells) and 35.6% to 48.0% (CD19-CAR iNK cells) (Fig. 2g). Subsequently, we monitored the changes in the dynamic ratio of CD19-CAR expression throughout the CD19-CAR iNK cell induction process. We observed comparable CD19-CAR expression ratios on day 7 (27.9% ± 3.4%, mean ± SD, n=3), Day 14 (27.2% ± 3.7%, mean ± SD, n=3), and day 21 (28.3% ± 2.2%, mean ± SD, n=3), with a slight increase on day 28 (33.4% ± 3.5%, mean ± SD, n=3) (Fig. 2j, k). Interestingly, CD19-CAR expressing cell ratios increased significantly on day 35 to 59.6% ± 2.9% (mean ± SD, n=3) and reached 62.3% ± 3.3% on day 42 (mean ± SD, n=3), which were comparable to expression ratios in engineered UCB-NK cells (61.3% ± 8.2%, mean ± SD, n=3) (Fig. 1h and Fig. 2j, k). To further ensure CD19-CAR expressing cell ratios of more than 90% in the final collection of CD19-CAR iNK cells on day 42, an additional step of enrichment of CD19-CAR^+^ CD34^+^ HSPCs on day seven during the HSPC expansion stage is required (Supplementary Figs. 2a-f).

Collectively, using our three-step induction strategy, a single CD34^+^ cell ultimately produced 1.4 × 10^7^ ± 0.1 × 10^7^ iNK cells and 7.6 × 10^6^ ± 1.2 × 10^6^ CD19-CAR iNK cells on day 42, and much higher yields of iNK cells (8.3 × 10^7^ ± 0.7 × 10^7^) and CD19-CAR iNK cells (3.2 × 10^7^ ± 0.2 × 10^7^) on day 49. A single umbilical cord blood unit of CD34^+^ cells (>1 × 10^6^) roughly has approximately the capacity to deliver 1.4 × 10^13^ - 8.3 × 10^13^ mature iNK or 7.6 × 10^12^ – 3.2 × 10^13^ CAR iNK cells, which prospectively ensures 760 to 83,000 doses (10^9^-10^10^ cells per dose) for treating patients.

### Molecular features and immune activities of CD34^+^ HSPC-derived iNK cells

The activation state and functionalities of NK cells are determined by the expression patterns of activating and inhibitory receptors (Fig. 3a). As expected, iNK cells expressed classical NK activating receptors, including CD319, NKp30, NKp44, NKG2D and CD69, and NK inhibitory receptors, including CD94, NKG2A and CD96^23, 24^. Furthermore, iNK cells also highly expressed critical NK effector molecules, including apoptosis-related ligands TRAIL and FasL^25^ (Fig. 3b). Subsequently, we analyzed CD107a^26^, a typical membrane protein associated with the cytotoxic activity of NK cells, along with tumor necrosis factor alpha (TNF-α)^27^, interferon gamma (IFN-γ), Perforin, and Granzyme B (GZMB)^28^. After exposure to PMA/ionomycin, CD107a, TNF-α, and IFN-γ were upregulated in UCB-NK and iNK cells. Phenotypic analysis revealed that both UCB-NK cells and iNK cells exhibited significant expression of cytotoxic granules, such as perforin and GZMB (Fig. 3c). Furthermore, statistical analysis did not show significant differences in protein expression patterns between UCB-NK cells and iNK cells (Fig. 3d).

**Fig.3.**
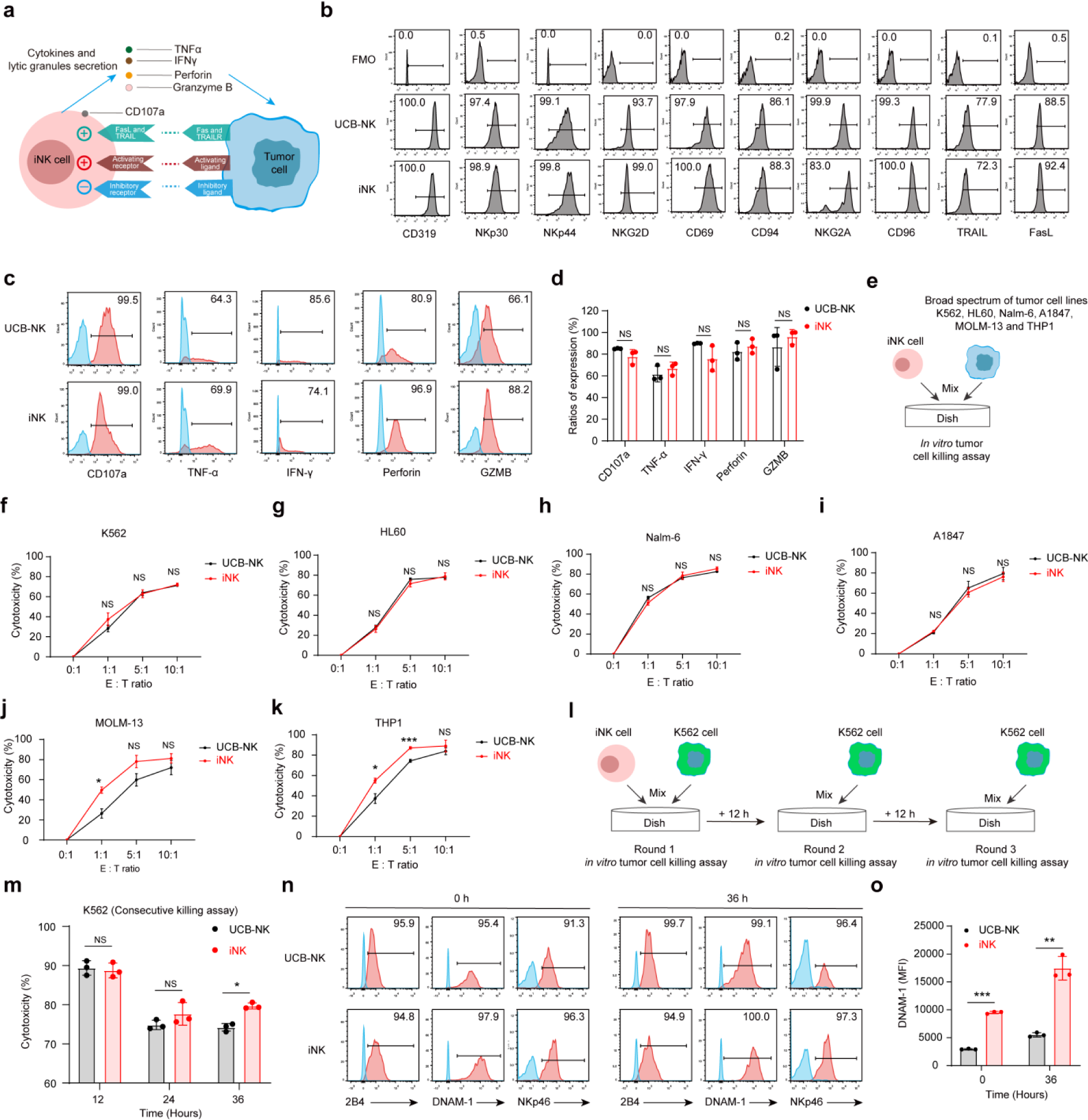
Molecular features and immune activities of CD34^+^ HSPC derived iNK cells. **a,** Schematic diagram showing the mechanisms of iNK cells that recognize and kill tumor cells. **b,** Flow cytometric analysis of typical NK receptors and effectors (CD319, NKp30, NKp44, NKG2D, CD69, CD94, NKG2A, CD96, TARIL, FasL). **c,** Representative flow cytometry histograms showing the expression levels of the CD107a, TNFα, IFN-γ, Perforin, and GZMB proteins in iNK cells and UCB-NK cells (CD45^+^CD56^+^). **d,** Statistical analysis of expression levels of CD107a, TNFα, IFN-γ, Perforin, and GZMB proteins (n = 3 repeats in each group). **e,** Schematic diagram of measuring the nonspecific cytotoxicities of iNK cells. **f-k,** Statistical analysis of nonspecific cytotoxicities of iNK cells and UCB-NK cells. (n = 3 repeats in each group). Cytotoxicity was calculated using a formula: (percentage of tumor cell death – percentage of tumor cell spontaneous death) / (1 - percentage of tumor cell spontaneous death) × 100. **l** Schematic diagram of measuring consecutive killing cytotoxicity of iNK cells. **m,** Statistical analysis of the consecutive cytotoxicity of iNK cells and UCB-NK cells. (n = 3 repeats in each group). **n** Representative flow cytometry histograms showing the expression levels of NK cell activation proteins (2B4, DNAM-1 and NKp46) before and after the consecutive killing assay in **m**. **o,** Statistical analysis of mean fluorescence intensity (MFI) of UCB-NK cells and iNK cells before and after the consecutive killing assay. (n = 3 repeats in each group). Data were presented as means ± SD. Independent two-tailed t test (**d**, **f-k**, **m** and **o**) or Mann-Whitney U test (**d** and **j**). NS, not significant, \**P* < 0.05, \*\**P* < 0.01, \*\*\**P* < 0.001.

Unlike adaptive T cells, NK cells possess an innate ability to recognize and eliminate tumor cells exhibiting reduced MHC-I expression^29^. Therefore, we first evaluated the tumoricidal capacity of iNK cells using various cancer cell lines, including the human erythroleukemia cell line K562, the human Caucasian promyelocytic leukemia cell line HL60, the human B lymphoblastic leukemia cell line Nalm-6, the human ovarian cancer cell line A1847, the human acute myeloid leukemia cell line MOLM-13, and the Epstein-Barr virus-negative B cell lymphoma cell line THP1 (Fig. 3e). The iNK cells showed similar tumor killing activities to those of UCB-NK cells at different E: T ratios (Fig. 3f-i). At lower E: In the T ratios, iNK cells exhibited superior cytotoxicity against MOLM-13 and THP1 (Fig. 3j,k). To evaluate the persistent cytotoxic activity of iNK cells, we performed three rounds of tumor-killing assays using K562 tumor cells at an effector-to-target (E: T) ratio of 1:1 (Fig 3l). iNK cells consistently demonstrated robust cytotoxicity, similar to the observation of UCB-NK cells in the first- and second-round killing assays. Intriguingly, iNK cells, but not UCB-NK cells, showed superior cytotoxicity in the third round killing assay (Fig. 3m, *P* < 0.05). Furthermore, we analyzed the expression patterns of the NK cell classical activation markers (2B4, DNAM-1, and NKp46) before and after three rounds of tumor cell killing tests (Fig. 3n)^30^The data revealed that both UCB-NK cells and iNK cells maintained high expression levels of these three activity receptors. It is noteworthy that iNK cells exhibited higher mean fluorescence intensities (MFI) for DNAM-1 regardless of the absence or presence of tumor cells (Fig. 3o). Collectively, CD34+ HSPC-derived iNK cells express typical markers related to NK cell function and possess the ability to kill broad-spectrum tumors.

### CD34^+^ HSPC-derived iNK cells suppressed tumor growth in A1847 tumor-bearing mice

To evaluate the *in vivo* therapeutic efficacy of HSPC-derived iNK cells, we established a human tumor cell line-derived xenograft model by intraperitoneal injection of luciferase-expressing A1847 cells (A1847-luci^+^, 2.0 × 10^5^ cells /mouse) into NCG mice (NOD/ShiLtJGpt-*Prkdc^em26Cd52^Il2rg^em26Cd^*^22^/Gpt strain) on day -1. We infused iNK cells (1.5 × 10^7^ cells / dose, i.p.) twice into the tumor-bearing animals on day 0 and day 7. In parallel, we used UCB-NK cells as a treatment control. Weekly bioluminescence imaging (BLI) was performed to capture the dynamics of tumor burden (Fig. 4a). As expected, iNK cells effectively suppressed tumor growth *in vivo*, demonstrating tumor-killing efficiencies comparable to those of UCB-NK cells (Fig. 4b, c). On the contrary, tumor-only animals exhibited progressive intensification of tumor burden, evidenced by increased radiance and total flux (Fig. 4b, c). iNK and UCB-NK cell-treated animals maintained stable body weight for 28 days. However, the tumor-only group showed significant body loss (Fig. 4d). We further investigated the dynamics of UCB-NK cells and iNK cells (CD45^+^CD56^+^) in the peritoneum after cell infusion (Fig. 4e). We detected 4.5% ± 4.0% (mean ± SD, n=4) of UCB-NK cells or 4.4% ± 3.2% (mean ± SD, n=4) of iNK cells in total nuclear cells of peritoneal dropsy from treated mice on day 7. As previously reported, CD56 expression decreased *in vivo* in tumor-bearing mice over time, indicating partially impaired activation of iNK cells (Fig. 4f)^31^. CD45^+^ CD56^+^ cells in the the peritoneum decreased to much lower levels in UCB-NK cell-treated mice (2.2% ± 1.7%, mean ± SD, n=4) and iNK cell-treated mice (1.6% ± 1.4%, mean ± SD, n=4) on day 14 (Fig. 4f, g). However, the kinetics of UCB-NK and iNK cells exhibited similar patterns (Fig. 4f, g). In particular, both the iNK cell and the UCB-NK cell significantly prolonged the survival of treated tumor-bearing animals (Tumor only: 35 days; Tumor + UCB-NK: 83 days; Tumor + iNK: 86 days; *P* < 0.001) (Fig. 4h). In conclusion, these results show that the CD34^+^ HSPC-derived iNK cells extend suppress tumor cell growth and markedly extend the survival of A1847-tumour-bearing animals.

**Fig.4.**
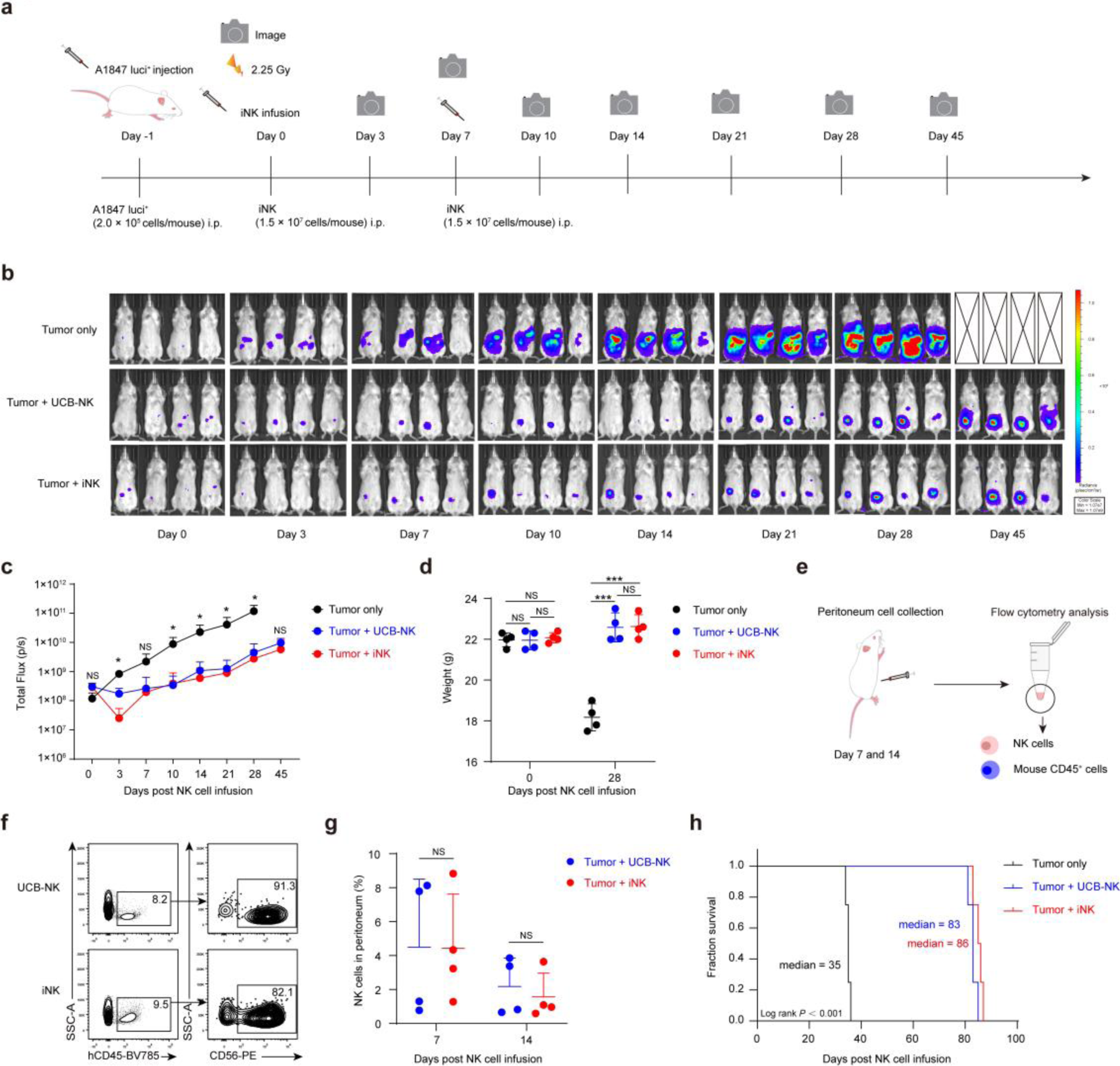
The iNK cell suppressed human tumor cell growth in xenograft animals. a, Schematic diagram of evaluation of nonspecific cytotoxicities of iNK cells in the tumor-bearing mouse. **b,** BLI analysis of the xenograft models (n = 4 in each group). **c,** Statistical analysis of total flux (photons/second, p/s) in xenograft models (n = 4 in each group). **d,** Statistical analysis of body weights of the xenograft models on day 28 after iNK cell injection (n = 4 in each group). **e,** Schematic diagram of analyzing the proportions of iNK or UCB-NK cells (CD45^+^CD56^+^) in NK cell-treated xenograft models. **f,** Representative FACS plots showing the proportions of iNK cells or UCB-NK cells (CD45^+^CD56^+^) in the peritoneum on day 7. **g,** Statistics analysis of the proportions of the iNK cells or UCB-NK cells (n = 4 in each group). **h,** Kaplan-Meier survival curves for xenograft models (*P* < 0.001, Log-rank test). Data were presented as means ± SD. One-way ANOVA test (**c**), Kruskal–Wallis tests (**c**), or two-tailed independent t test (**c**, **d,** and **g**). NS, not significant, \**P* < 0.05, \*\*\**P* < 0.001.

### CD19-CAR iNK cells efficiently eliminated B cell lymphoma and leukemia tumor cells

To confirm the specific cytotoxicity of CD19-CAR iNK cells, we performed *in vitro* tumor-killing assays by coculture of CD19-CAR iNK cells with CD19-positive Nalm-6 cells, primary tumor cells of CD19-positive human B cell lymphoma and leukemia, respectively. We first selected the Nalm-6 tumor cell line for the tumor-killing effect of CD19-CAR iNK cells. Initially, Nalm-6 cells (Targets, T) were incubated with UCB-NK cells, iNK cells, and CD19-CAR iNK cells (Effectors, E) respectively at E: T = 0:1, 0.2:1, 0.4:1, 0.8:1, 1.6:1, and 5: 1 for 4 hours (Fig. 5a). As expected, CD19-CAR iNK cells showed superior cytotoxicity against tumor targets than UCB-NK cells and iNK cells (Fig. 5b, *P* < 0.001). We then analyzed the expression of CD107a, a typical membrane protein associated with NK cell cytotoxicity^26^. Our result showed that CD19-CAR iNK cells exhibited much higher levels of CD107a expression than UCB-NK cells and iNK cells (Fig. 5c, d, *P* < 0.001). Furthermore, we also evaluated the nonspecific tumoricidal capacity of CD19-CAR iNK cells using K562 tumor cells (Fig. 5e). Our results indeed demonstrated that these CD19-CAR iNK cells retained nonspecific cytotoxicity similar to that of UCB-NK and iNK cells (Fig. 5f). To assess the sustained cytotoxic activity of CD19-CAR iNK cells, we performed three rounds of tumor-killing assays with Nalm-6 tumor cells at an effector-to-target (E: T) ratio of 1:1 (Fig. 5g). Throughout three rounds, CD19-CAR iNK cells maintained constant cytotoxicity. It is noteworthy that the serial killing feature was only observed in CAR iNK cells but not in UCB-NK and iNK cells (Fig. 5h, i, *P* < 0.001). Therefore, CD19-CAR HSPC-derived CD19-CAR iNK cells possess specific cytotoxicity against CD19 positive tumor cells and maintain the nonspecific tumoricidal capacity of NK cells. Next, we tested the particular cytotoxicity of CD19-CAR iNK against CD19-positive primary tumor cells from patients with B cell lymphoma and B cell leukemia (Fig. 5j). Our results showed that CD19-CAR iNK cells exhibited increased cytotoxicity against patient-derived B cell lymphoma (Patient 1) and B cell leukemia cells (Patient 2 and 3) (*P* < 0.001). However, iNK cells showed limited cytotoxicity against these three patient tumor cells even at higher E: T ratios (Fig. 5k-n).

**Fig.5.**
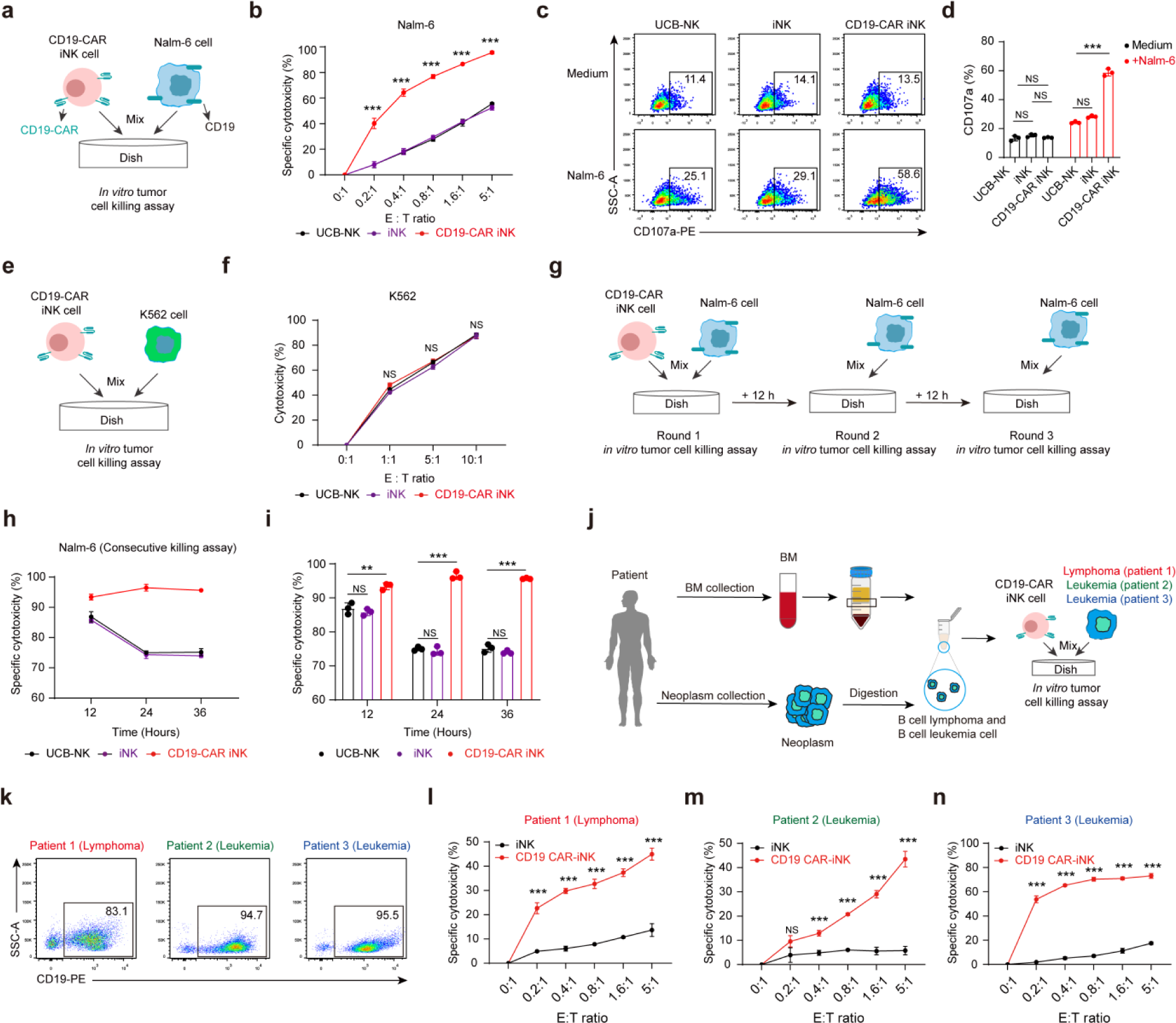
CD19-CAR iNK cells exhibited specific tumor-killing activities against B-cell lymphoma and leukemia *in vitro*. **a,** Schematic diagram of evaluation of the *in vitro* specific cytotoxicities of CD19-CAR iNK cells. **b,** Statistic analysis of the specific cytotoxicities of CD19-CAR iNK cells. **c,** Representative FACS plots showing the expression levels of the CD107a protein of CD19-CAR-iNK cells. **d,** Statistical analysis of the expression levels of the CD107a protein (n = 3 repeats in each group). **e,** Schematic diagram of evaluation of nonspecific cytotoxicities of CD19-CAR iNK cells. **f,** Statistical analysis of the nonspecific cytotoxicities of CD19-CAR iNK cells. (n = 3 repeats in each group). **g,** Schematic diagram of the evaluation of the consecutive killing cytotoxicities of CD19-CAR iNK cells. **h-i,** Statistical analysis of consecutive specific cytotoxicities of CD19-CAR iNK cells. (n = 3 repeats in each group). **j,** Schematic diagram showing the specific cytotoxicities of CD19-CAR iNK cells against CD19-expressing tumor cells isolated from patients with B cell leukemia or B cell lymphoma. BM, bone marrow. **k,** Representative FACS plots showing the expression levels of the CD19 protein in patient-derived B-cell lymphoma or leukemia cells (Gated from CD45^+^). **l,** Statistical analysis of the specific cytotoxicities of CD19-CAR iNK cells against patient-derived B cell lymphoma cells. (n = 3 repeats in each group). **m-n,** Statistical analysis of the specific cytotoxicities of CD19-CAR iNK cells against patient-derived B cell leukemia cells. (n = 3 for each group). Cytotoxicity was calculated using the formula: (percentage of tumor cell death – percentage of tumor cell spontaneous death) / (1 – percentage of tumor cell spontaneous death) × 100. Data were presented as means ± SD. One-way ANOVA test (**b** and **f**) or two-tailed independent t test (**d**, **i**, **l**, **m** and **n**). NS, not significant, \*\**P* < 0.01, \*\*\**P* < 0.001.

Collectively, CD19-CAR iNK cells exhibit superior specific cytotoxicity against CD19-positive tumor cells and still possess serial killing capacity and maintain nonspecific cytotoxicity of NK cells.

### CD19-CAR iNK cells suppressed tumor growth in Nalm6 tumor-bearing mice

To evaluate the therapeutic efficacy *in vivo* of CD19-CAR iNK cells, we established B-ALL xenograft animal models using Nalm-6 cells expressing luciferase (Nalm-6 luci^+^). On day 1, B-NDG hIL15 mice (NOD.CB17-*Prkdc^scid^Il2rg^tm1^Il15^tm^*^1^(IL15)/Bcgen background) were injected intravenously with 1 × 10^5^ Nalm-6 luci^+^ cells. Tumor-bearing mice were injected with control iNK cells or CD19-CAR iNK cells (the equivalent of 1.0 × 10^7^ CD19-CAR iNK cells/dose, twice) through the tail veins. Tumor progression was monitored weekly using bioluminescent imaging (BLI) (Fig. 6a). iNK cell treatment alone did not suppress Nalm-6 cell growth *in vivo*, consistent with the observations in UCB-NK cell-treated Nalm-6 tumor-bearing animals^22^. As expected, CD19-CAR iNK cells effectively suppressed tumor growth *in vivo*, demonstrating superior tumor-killing efficiencies over iNK cells (Fig. 6b, c). We further investigated the *in vivo* kinetics of iNK cells and CD19-CAR iNK cells in tumor-bearing mice (Fig. 6d). We observed apparent human CD45^+^CD56^+^ iNK cells that circulate in the peripheral blood of the animals treated with iNK cells (7.4% ± 2.2%, mean ± SD, n=5)- and iNK cells treated with CD19-CAR (10.0% ± 1.7%, mean ± SD, n=5)-treated animals on day 7. However, circulated CD45^+^CD56^+^ decreased too much lower levels in mice treated with iNK cells (1.2% ± 0.8%, mean ± SD, n=5) and mice treated with CD19-CAR iNK cells (1.9% ± 1.4%, mean ± SD, n=5) on day 14 (Fig. 6e, f). In particular, the CD19-CAR iNK cell therapy significantly extended the survival of Nalm-6 tumor-bearing mice (Tumor only: 25 days; Tumor + iNK: 27 days; Tumor + CD19-CAR iNK: 49 days; *P* < 0.001) (Fig. 6g).

**Fig.6.**
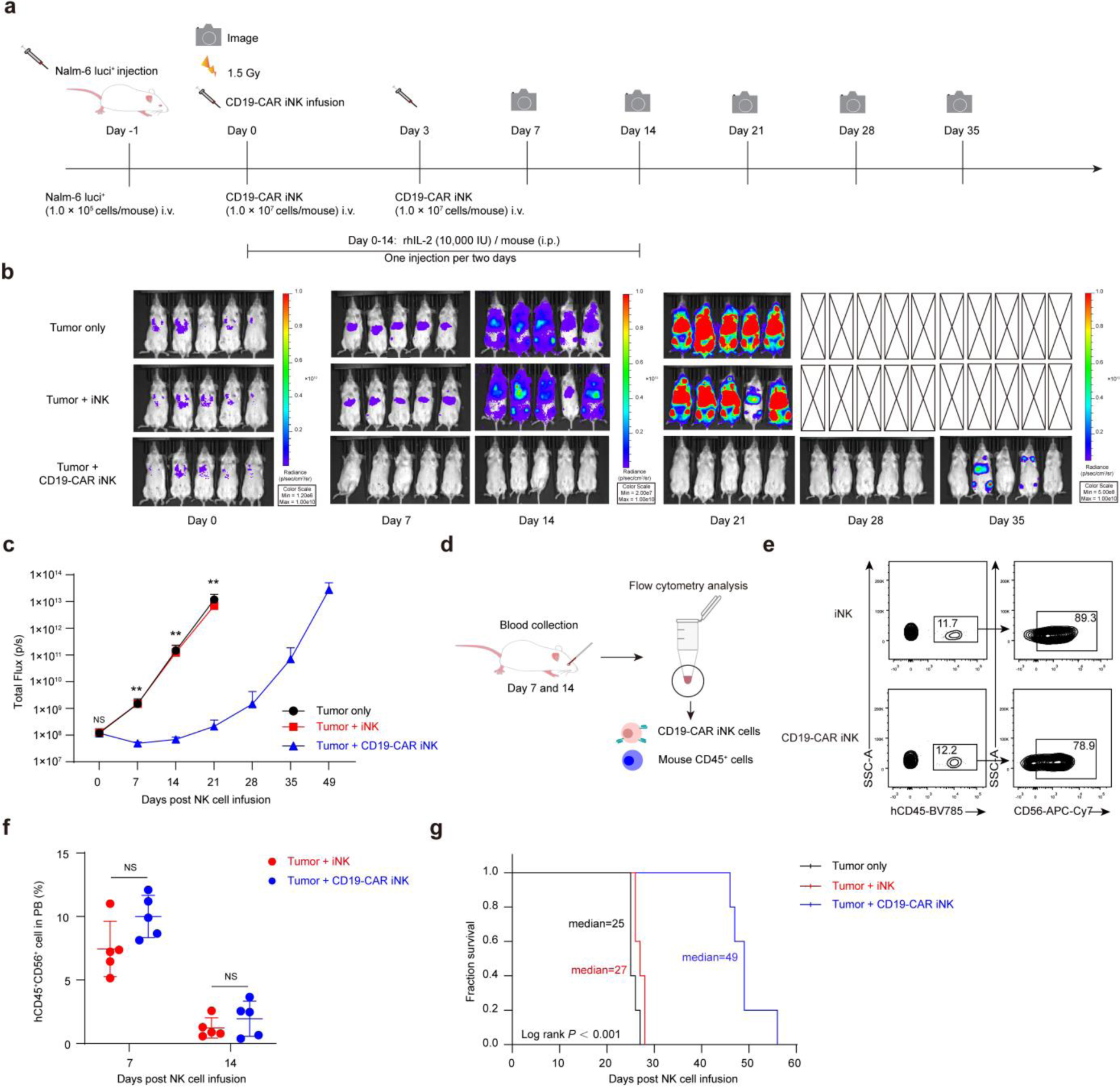
The CD19-CAR iNK cell suppressed the growth of human B leukemia cells in xenograft animals. **a,** Schematic diagram of the evaluation of the cytotoxicities of CD19-CAR iNK cells *in vivo*. **b,** BLI analysis of the xenograft models (n = 5 in each group). **c,** Statistical analysis of total flux (photons/second, p/s) in xenograft models (n = 5 in each group). **d,** Schematic diagram for analyzing the proportion of human CD45^+^CD56^+^ cells in peripheral blood (PB). **e,** Representative FACS plots showing the proportion of human CD45^+^CD56^+^ cells from PB in tumor-bearing mice on day 7. **f,** Statistical analysis of the ratios of CD45^+^ CD56^+^ peripheral blood NK cells in tumor-bearing mice on day 7 and day 14 (n = 5 in each group). **g,** Kaplan-Meier survival curves for xenograft models (*P* < 0.001, Log-rank test). Data are presented as means ± SD. One-way ANOVA test (**c**), Kruskal–Wallis tests (c), or two-tailed independent t-test (**f**). NS, not significant, \*\**P* < 0.01.

Our data demonstrate that CD19-CAR iNK cells can efficiently kill CD19-positive tumor cells *in vivo* and prolong the survival of tumor-bearing animals.

### Cryopreserved CD19-CAR iNK cells retained tumor-killing efficacies *in vitro* and *in vivo*

To mimic the potential ‘off-the-shelf’ products of CD19-CAR iNK cells, we harvested and cryopreserved CD19-CAR iNK cells using commercial GMP grade CryoStor® CS10 cell freezing medium^32^. After six months, cryopreserved cells were thawed and revived for recovery tests for 24, 48, and 72 hours to assess the dynamic variations of their viability, cell numbers, and tumor-killing activities (Fig. 7a). After 72 hours of *in vitro* revival, the viabilities of NK cells reached 87.8% ± 1.1% (mean ± SD, n = 3). The CD19-CAR iNK cell count was 90.0% ± 1.1% (mean ± SD, n = 3) (Fig. 7b, c).

**Fig.7.**
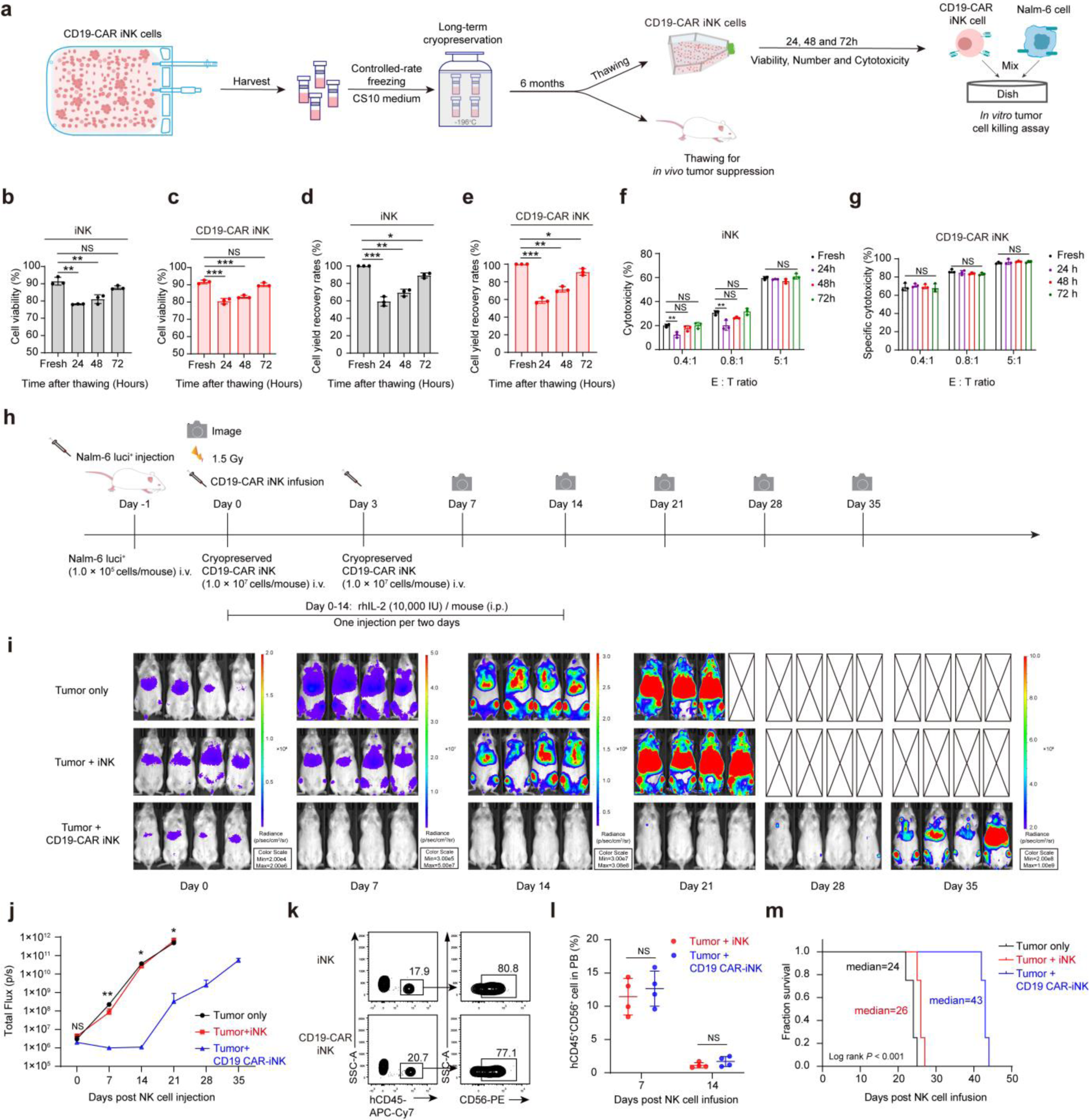
The cryopreserved CD19-CAR iNK cells retained tumor-killing efficacies *in vitro* and *in vivo*. **a,** Schematic diagram for evaluating the specific cytotoxicities of the cryopreserved CD19-CAR iNK cells *in vitro* and *in vivo*. **b-c,** Statistical analysis of the viabilities of iNK cells and CD19-CAR iNK cells after thawing for 24 hours, 48 hours, and 72 hours. (n = 3 repeats in each group). **d-e,** Statistical analysis of cell yield recovery rates of iNK cells and CD19-CAR iNK cells after thawing for 24 hours, 48 hours, and 72 hours. (n = 3 for each group). **f-g,** Statistical analysis of the specific cytotoxicities of iNK cells and CD19-CAR iNK cells after thawing for 24 hours, 48 hours, and 72 hours. (n = 3 repeats in each group). **h,** Schematic diagram of the evaluation of the specific cytotoxicities of the cryopreserved CD19-CAR iNK cells *in vivo*. **i,** BLI analysis of the xenograft models (n = 4 in each group). **j,** Statistical analysis of total flux (photons/second, p/s) in xenograft models (n = 4 in each group). **k,** Representative FACS plots showing the proportion of iNK cells and CD19-CAR iNK cells (CD45^+^CD56^+^) from peripheral blood on day 7. **l,** Statistics analysis of the ratios of iNK cells and CD19-CAR iNK cells (CD45^+^CD56^+^) from peripheral blood on day 7 and day 14 (n = 4 in each group). **m,** Kaplan-Meier survival curves for xenograft models (*P* < 0.001, Log-rank test). Cytotoxicity was calculated using the formula: (percentage of tumor cell death – percentage of tumor cell spontaneous death) / (1 – percentage of tumor cell spontaneous death) × 100. Data were presented as means ± SD. Independent two-tailed t test (**b**, **c**, **d**, **e**, **f,** and **l**), one-way ANOVA test (**g** and **j**), or Kruskal–Wallis tests (**j**). NS, not significant, \**P* < 0.05, \*\**P* < 0.01, \*\*\**P* < 0.001.

Furthermore, after 72 hours of *in vitro* culture, the iNK cell and CD19-CAR iNK cell counts were restored to 89.2% ± 2.8% (mean ± SD, n = 3) and 91.2% ± 3.8% (mean ± SD, n = 3) of pre-cryo-preservation input counts, respectively (Fig. 7d, e). Furthermore, Nalm-6 tumor-killing assays showed that revived CD19-CAR iNK cells exhibited similar cytotoxicities to fresh CD19-CAR iNK cells (Fig. 7f, g).

Furthermore, we evaluate the *in vivo* tumor killing efficacy of cryopreserved CD19-CAR iNK cells (Fig. 7h). As expected, thawed CD19-CAR iNK cells without extended culture revival still retained enhanced antitumor activity over NK cells *in vivo* (Fig. 7i, j). We also observed apparent human CD45^+^CD56^+^ iNK cells circulating in the peripheral blood of the animals treated with iNK cells (11.4% ± 2.8%, mean ± SD, n=4)- and CD19-CAR iNK cells (12.7% ± 2.6%, mean ± SD, n=4)-treated animals on day 7. However, circulating CD45^+^CD56^+^ decreased too much lower levels in mice treated with iNK cells (1.1% ± 0.4%, mean ± SD, n=4) and mice treated with CD19-CAR iNK cells (1.7% ± 0.7%, mean ± SD, n=5) on day 14 (Fig. 7k, l). In particular, thawed CD19-CAR iNK cell therapy significantly extended the survival of Nalm-6 tumor-bearing mice (Tumor only: 24 days; Tumor + iNK: 26 days; Tumor + CD19-CAR iNK: 43 days; *P* < 0.001) (Fig. 7m).

In summary, our data demonstrate that cryopreserved CD19-CAR iNK cells maintain antitumor activity both *in vitro* and *in vivo*.

## Discussion

In this study, we develop an efficient three-step strategy that can generate trillions of iNK cells and CD19-CAR iNK cells from a single umbilical cord blood unit of CD34^+^ HSPCs. The generating efficiencies of iNK cells and CD19-CAR iNK cells are 6,918-20,224 times over traditional methods^10^. Interestingly, our method sharply reduces the engineering cost of CD19-CAR to negligible. iNK cells derived from CD34^+^ HSPC and CD19-CAR iNK cells possess ideal tumoricidal activities against human tumors. Our study provides profound insight into the use of CD34^+^ HSPCs as cell sources to generate CAR NK cells and expand their accessibility and affordability for patients.

Our CD19-CAR iNK cells showed CD19-CAR positivity rates ranging from 58.5% to 64.4%, comparable to the positivity rates of CD19-CAR NK cells obtained by engineering mature UCB-NK cells (47.8% to 87.4%, median = 66.6%)^33^. Transduction of CAR into CD34^+^ HSPCs does not alter the differentiation efficiencies of the generation of CAR iNK cells or results in silencing of CAR expression in derived iNK cells. Furthermore, in a mouse model with Nalm-6 xenograft, CD19-CAR iNK cells showed superior specific killing activity compared to unmodified iNK cells, consistent with previous studies in which cryopreserved and thawed CD19-CAR NK cells (injected with 5 × 10^6^ CD19-CAR NK cells per dose, four times) without any culture revival significantly inhibited the growth of Nalm-6 tumors in mice^34^. By administering 1 × 10^7^ CD19-CAR iNK cells twice, we achieved similar suppression effects on tumor growth, confirming the therapeutic potential and application value of CD19-CAR iNK cells in antitumor therapy. However, unlike long-term therapeutic efficacies observed via single-dose CD19 CAR-T treatment in animal models and in certain patients, two doses of CD19-CAR iNK cell treatment still lack persistent therapeutic efficacies in tumor-bearing animals treated, indicating that multiple doses and high doses of CAR-iNK cell therapy are critical for persistent suppression of tumors. However, our method produces reliable large-scale NK and CAR NK cells from CD34^+^ HSPCs to prospectively treat human cancers.

Over decades, the AFT024 cell line has been shown to be a feeder cell priority for the expansion of human CD34^+^ HSPCs, which ideally maintains the stemness and curbs the differentiation of human long-term hematopoietic stem cells (HSCs)^17–19^. In our approach, AFT024 cells further expanded CD34^+^ HSPCs without losing their differentiation potential from the NK lineage. The 14-day expansion based on AFT024 feeder cell-based 14-day expansion in the presence of HSC culture medium^20^ significantly increases the iNK and CAR iNK cell yields more than 160 times, indicating that the potential of the NK lineage is preserved primarily in expanded CD34^+^ HSPCs. Our method reliably produces large-scale homogeneous iNK cells and CAR iNK cells from a single donor umbilical cord blood unit of CD34^+^ HSPCs, which prospectively ensures the need for 760 to 83,000 doses (10^9^ - 10^10^ cells per dose)^8, 9^ to treat tumor patients. Our result shows that the thawed CD19-CAR iNK cells retain anti-tumor efficacy, providing obvious leverages for long-term safety inspections of CAR NK products and comprehensive clinical applications. Thus, in clinical settings, the tumorigenic risks of CAR iNK cells can be well evaluated in animal models before large-scale clinical applications, as autogenous CAR-T cell therapies reportedly show a few cases of CAR-T-derived secondary T cell tumors^35^. Our method shows that engineering CAR at the CD34^+^ HSPC stage barely harms iNK induction efficiencies. Consequently, we have reduced the use of CAR engineering materials to 1 / 140,000 –1 / 600,000, the resulting cost of which is negligible for the manufacturing of CAR NK cells. Interestingly, the CD19-CAR expression rates in our method maintain consistently high levels (58.5% to 64.4%), indicating that there is no obvious silencing of CAR expression during the entire three-step process. Thus, reducing the manufacturing cost of CAR iNK cells will promote the accessibility of this type of cell therapy.

In conclusion, we have developed a comprehensive technique for efficiently generating massive CD34^+^ HSPC-derived iNK cells and CAR iNK cells, with characteristics of trillion-scale yields and negligible cost of CAR engineering from a single donor umbilical cord blood unit of CD34^+^ HSPCs. Our study provides profound insight into the use of CD34^+^ HSPCs as cell sources to generate CAR NK cells and expand their accessibility and affordability for patients.

## Materials and Methods

### Ethics statement

NCG mice and B-NDG hIL15 mice were housed in SPF-grade animal facilities at the Guangzhou Institutes of Biomedicine and Health, Chinese Academy of Sciences. All animal-related procedures in this study received approval from the Institutional Animal Care and Use Committee of the Guangzhou Institutes of Biomedicine and Health. NK cell antitumor activity assessments in animals received approval from the Biomedical Research Ethics Committee of the Guangzhou Institutes of Biomedicine and Health, Chinese Academy of Sciences. The use of patient samples was carried out in accordance with the provisions of the Declaration of Helsinki. All patient samples were collected with prior consent signatures from the patient and reviewed and approved by the ethics committee of the Jinan University First Affiliated Hospital. This informed consent included allowing patients to publish the data arising from the tissues they obtained.

### Statistics

All quantitative analyses were performed with SPSS (version 23, IBM Corp., Armonk, NY, USA). The Shapiro-Wilk normality test in SPSS was used to evaluate the normal distribution of the data. The two-tailed independent t test (for normally distributed data) and the Mann-Whitney U test (for nonnormally distributed data) were applied for comparison of two groups of data. For three or more groups: the one-way ANOVA test (for normally distributed data) and Kruskal-Wallis tests (for nonnormally distributed data) were used. Survival curves were plotted using the Kaplan-Meier method. Differences in survival rates between groups were evaluated using the Log rank test (Mantel-Cox). Statistical analyses were performed using GraphPad Prism (8.0.2, GraphPad Software).

## Data availability

Single-cell RNA sequencing data (fastq files) have been uploaded to the public database of the Genome Sequence Archive (HRA007978). Raw flow cytometry data files and bioluminescent imaging data are available upon request. Other relevant information or data are available from the corresponding authors upon reasonable request.

## Acknowledgements

This work was supported by grants from the National Natural Science Foundation of China (81925002) and the National Key R&D Program of China (2020YFA0112404).

## Author contributions

J.L., Y.W., and X.Z. performed the core experiments and contributed equally to this work. J.L. wrote the manuscript. Y.L. and Q.W. analyzed the RNA-seq data. X.L., Y.G., H.W., L.L., H.P., B.W., D.H., C.X., T.W., and M.Z. participated in multiple experiments. X.D., H.Z., F.D., Y.Z. and X.Z. discussed the data and manuscript. J.W. and F.H. designed the project and wrote the manuscript. J.W. provided the competing interests.

---

**Supplementary Fig.1.**
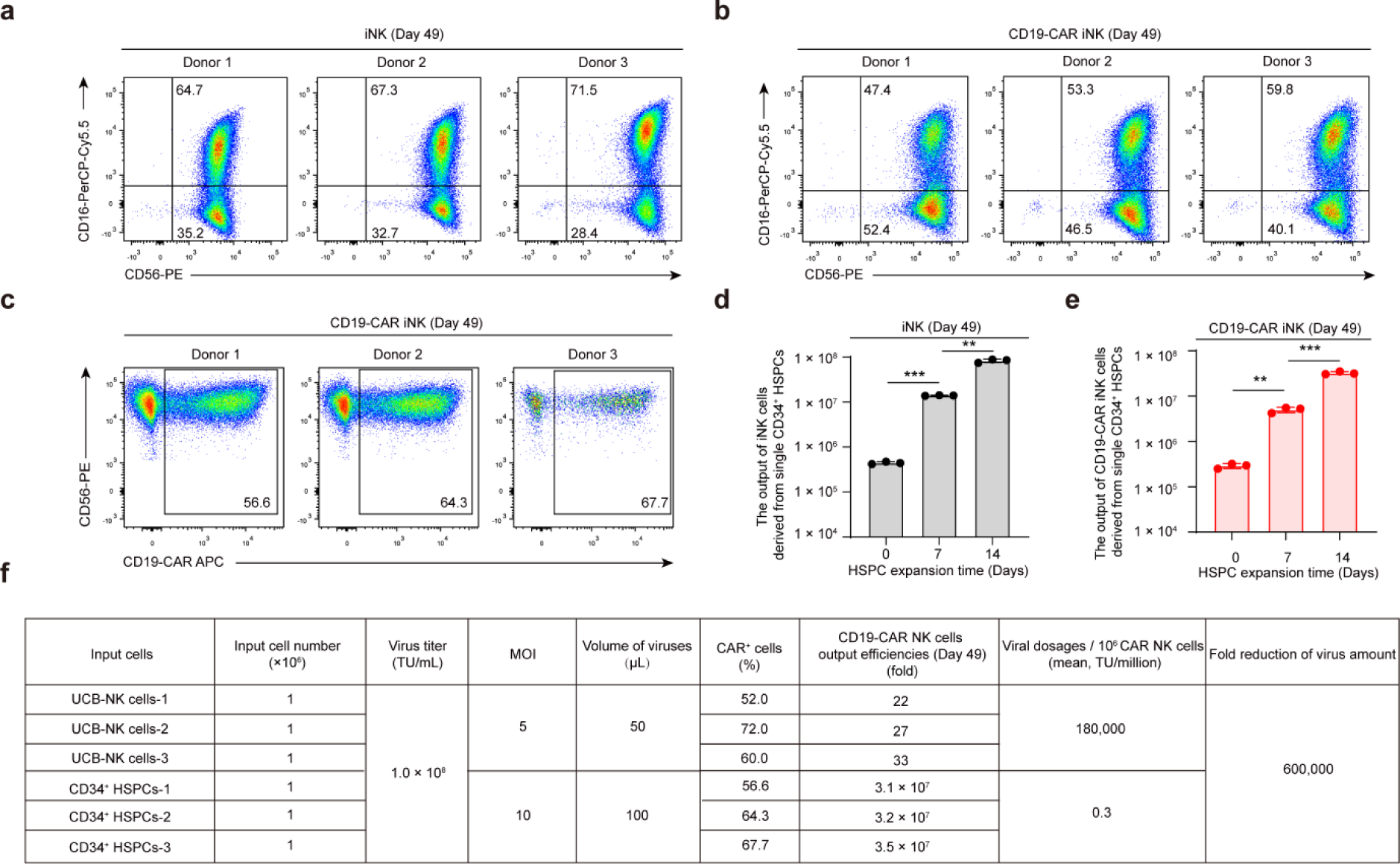
Immune phenotypes and yields of iNK cells and CD19-CAR iNK cells after 3-week bag-based culture. **a-b,** flow cytometric analysis of the immune phenotypes of iNK cells (**a**) and CD19-CAR iNK cells (**b**) (CD56^+^ CD16^+/-^). Data were collected from three donor umbilical cord blood units. **c,** Flow cytometric analysis of CD19-CAR expression (CD56^+^CD19-CAR^+^). **d-e,** Statistical analysis of the output efficiencies of iNK cells (CD45^+^CD3^-^CD56^+^CD16^+/-^) or CD19-CAR iNK cells (CD45^+^CD3^-^CD56^+^CD16^+/-^CD19-CAR^+^) derived from single CD34^+^ HSPCs or CD19-CAR HSPCs. Day 0, fresh CD34^+^ HSPCs. Day 7, 7-day expanded CD34^+^ HSPCs. Day 14, 14-day expanded CD34^+^ HSPCs. **f**, Table showing the calculated quantities of CD19-CAR retroviruses (TU). The viral particles required to generate 1.0 × 10^6^ CD19-CAR iNK cells and CD19-CAR NK cells were compared. Data were presented as means ± SD. Independent two-tailed t test (**d** and **e**). NS, not significant, \*\**P* < 0.01, \*\*\**P* < 0.001.

**Supplementary Fig.2.**
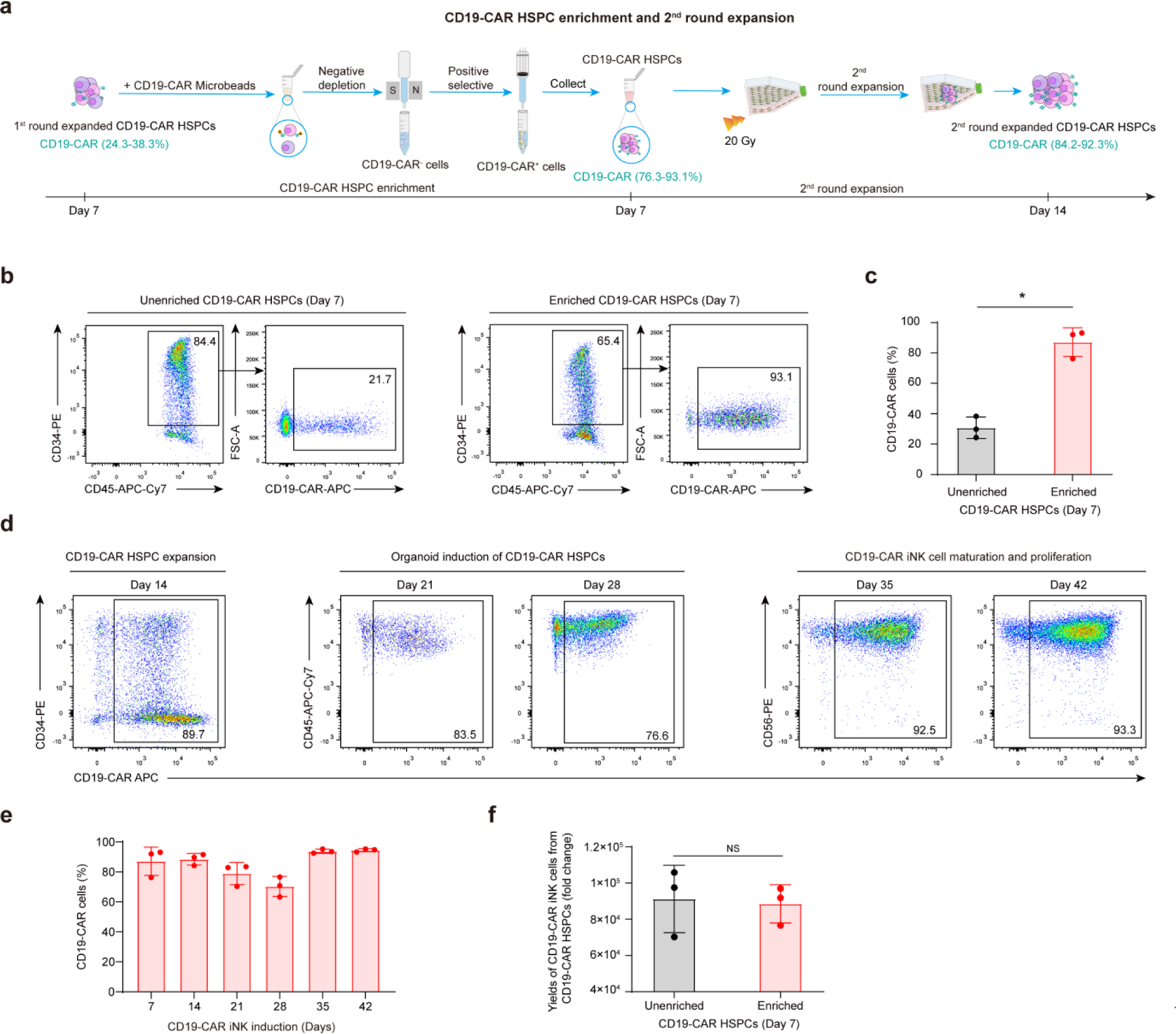
Enrichment of CD19-CAR^+^ CD34^+^ HSPCs on day 7 further increased the positive CD19-CAR ratios of CD19-CAR iNK cells on day 42 by more than 90%. **a,** Schematic diagram of the generation of CD19-CAR^+^ HSPC cells from expanded 7-day CD19-CAR HSPCs. **b,** Flow cytometric analysis of the CD19-CAR expression in CD34^+^ HSPCs before and after enrichment using a CD19-CAR based microbead enrichment process on Day 7. **c,** Statistical analysis of CD19-CAR expression levels in CD34^+^ HSPC before and after enrichment. Data were collected from three donor umbilical cord blood units. **d,** Representative FACS plots showing the expression of CD19-CAR of total cells at the indicated time points. **e,** Statistical analysis of the CD45^+^ CD19-CAR^+^ cell ratios at the indicated time points. (n = 3 in each group, n indicated three donor umbilical cord blood units). **f**, Statistical analysis of the yields of CD19-CAR iNK cells from single 7-day expanded CD19-CAR HSPCs. Data were presented as means ± SD. Independent two-tailed t test (**c** and **f**). NS, not significant, \**P* < 0.05.

